# Estradiol alters actin and protrusion dynamics in endometriotic epithelial cells

**DOI:** 10.1101/2025.05.14.654086

**Authors:** Shohini Banerjee, Corey Herr, Wolfgang Losert, Kimberly M. Stroka

**Affiliations:** Fischell Department of Bioengineering, University of Maryland, College Park, MD, USA; Department of Physics, University of Maryland, College Park, MD, USA; Institute for Physical Science and Technology, University of Maryland, College Park, MD, USA; Marlene and Stewart Greenebaum Comprehensive Cancer Center, University of Maryland, Baltimore, MD, USA; Biophysics Program, University of Maryland, College Park, MD, USA; Center for Stem Cell Biology and Regenerative Medicine, University of Maryland, Baltimore, MD, USA

## Abstract

Estradiol (E2), a sex steroid hormone molecule, plays a key role in regulating the actin and shape dynamics of cells in a multitude of normal and pathophysiological conditions. While cytoskeletal rearrangements, membrane dynamics, and cellular protrusions are intimately involved in cell motility and invasiveness, little is known about the impact of E2 on these processes in estrogen-dependent epithelial cells. In this study, we quantified the impact of E2 on cell shape and actin dynamics in 12Z human endometriotic epithelial cells transfected with LifeAct-GFP and observed with lattice lightsheet microscopy, a new imaging technique fast enough to capture 3D dynamics on second timescales. E2, when applied for 24 hours, significantly decreased cell circularity, solidity, and rate of change of circularity, indicating a transition to a more elongated and less variable morphology. 24-hour E2 treatment also induced the formation of large membrane protrusions reminiscent of invadopodia and led to a more disordered flow of actin within those protrusions. However, these effects were not seen after 15 minutes of E2 treatment, suggesting that longer-term signaling is required to drive these structural changes. Together, these results suggest that E2 modulates actin polymerization and membrane protrusion dynamics in endometriotic epithelial cells and may prime them for cell invasion. This work highlights a role for hormonal signaling in mediating cytoskeletal plasticity and migratory cell phenotypes.

## I. INTRODUCTION

The ability of cells to dynamically alter their structure and shape plays a critical role in numerous healthy and disease processes, such as cell proliferation, migration, invasion, and tissue remodeling. Estrogens can play a key role in regulating these cellular activities. For example, estrogens are known to mediate cell invasiveness in estrogen-dependent conditions, such as endometriosis [1–4], estrogen receptor-positive breast cancers [5–8], and gynecological cancers [9– 12]. Moreover, estrogens have been linked to cytoskeletal remodeling [1,13–15], which likely influences cell motility and morphodynamic flexibility – a trait which helps cells to migrate through diverse environments such as tissue microtracks and peritoneal, vascular, and lymphatic barriers.

Actin dynamics play a key role in directed cell migration [16–18] and the formation of cellular protrusions such as filopodia [19], lamellipodia [20], and invadopodia [21,22]. Upregulating actin polymerization by itself does not create protrusions, however changing actin cytoskeleton dynamics can drive formation and suppression of protrusions [23]. The impact of estrogen on protrusion dynamics and cytoskeletal arrangement over time remains unclear. Moreover, most previous cytoskeleton studies fix cells prior to immunofluorescence, which does not allow for the observation of real-time, live-cell dynamics. To overcome this, actin dynamics have been studied in mammalian cells transfected with GFP-tagged actin and imaged using methods such as 2D fluorescence microscopy [24] and the nanoscale precise imaging by rapid beam oscillation (nSPIRO) method [25] for 3D, but these techniques are still limited in terms of resolution and photobleaching.

In the present study, we sought to clarify the effect of 17β-estradiol (E2), a dominant and potent estrogen molecule, on cytoskeletal and shape dynamics in the 12Z human endometriotic epithelial cell line. This cell line has been designed to model estrogen-dependent epithelial behavior, as seen in endometriosis [26]. We used 3D lattice light sheet microscopy to capture unprecedented volumetric time-lapses of LifeAct-GFP-labeled actin in live 12Z cells. Live-cell imaging revealed dynamic cytoskeletal reorganization, 3D membrane protrusions, and membrane ruffling. We quantified the effects of E2 on cell shape, membrane protrusion dynamics, and actin polymerization using custom image analysis pipelines, including optical flow analysis. Our study reveals that 24-hour E2 exposure significantly alters cell morphodynamic behavior, increases protrusiveness, and disrupts actin polymerization coordination in 12Z cells. These findings provide new insight into the role of estrogen in modulating epithelial structural plasticity and promoting invasive phenotypes in estrogen-dependent diseases.

## II. METHODS

### A. Cell culture

The 12Z cell line was purchased from Applied Biological Materials (Cat. #T0764). Cells were cultured in DMEM/F12 (Gibco) supplemented with 10% FBS (Gibco) and 1% penicillin-streptomycin (Gibco). For estradiol studies, about 48 hours prior to imaging, the culture media was replaced with phenol-free DMEM/F12 (Gibco) supplemented with 10% charcoal-stripped FBS (Gibco) and 1% penicillin-streptomycin (Gibco). 17β-estradiol (E2) was purchased from Sigma Aldrich (Cat. #E8875-1G) in powder form and dissolved in 100% ethanol to create a stock solution. Cells were treated with a vehicle control (0.001% ethanol) or 10 nM E2 for 15 minutes or 24 hours prior to imaging. This E2 concentration and incubation time was chosen according to prior literature [27,1,3,8]. The cell line was authenticated by the company prior to shipping and our experiments were conducted at passages 58-68.

### B. Cell transfection and staining

The LifeAct-GFP plasmid, a probe for globular and filamentous actin, was a generous gift from Dr. Denis Wirtz’s lab (Johns Hopkins University, MD, US). 12Z cells were transfected with a LifeAct-GFP probe for globular and filamentous actin using electroporation technology. Briefly, cells were suspended in 100µL of a supplemented nucleofector solution (Lonza, Cell Line Nucleofector Kit L, cat. #VCA-1005, Germany) with 5 µg of LifeAct-GFP plasmid, transferred to a cuvette, and subjected to electroporation via the Nucleofector device (Amaxa, Germany). The Nucleofector program X-001 was selected based on viability optimization experiments. Transfected cells were immediately seeded on a chambered coverslip (Ibidi, Germany) coated with 10 µg/mL collagen I from rat tail (Corning, cat. #354249, NY, US) and incubated overnight prior to imaging. For validation studies, to ensure that the borders of the actin cytoskeleton were coincident with the plasma membrane, cells were exposed to a plasma membrane stain (CellMask, ThermoFisher, cat. #C56129, MA, US).

### C. Lattice lightsheet microscopy

Fluorescent actin time-lapses were captured using the Zeiss Lattice Light Sheet 7 (LLS7) microscope equipped with the Hamamatsu ORCA-Fusion camera system and the ZEN Microscopy software (Zeiss, Germany). The 30 µm x 1000 nm light sheet was utilized for all imaging. LifeAct-GFP was visualized using the 488 nm laser and for cells with the additional plasma membrane stain, the 560 nm laser was also used. Image slices were 0.2 µm in thickness. Cells were maintained at 37°C in 90% humidity and 5% CO_2_ for the duration of imaging.

### D. Shape dynamics analysis

Cell morphodynamics were analyzed from fluorescence maximum intensity projections obtained from lattice lightsheet microscopy. Time-lapse image stacks were loaded into ImageJ and individually thresholded for optimal object segmentation. Morphology parameters, including circularity, aspect ratio, and solidity, were then obtained from binarized image stacks. Circularity is defined by the formula 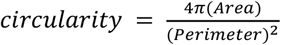, where a value of 1.0 indicates a perfect circle and a value closer to 0 indicates an elongated polygon. Aspect ratio is defined as 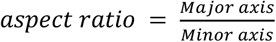. Solidity is defined by 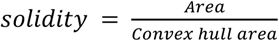, where a cell that is perfectly circular would have a value of 1.0 and a cell that has protrusions and/or indentations would have a value closer to 0. We refer to the absolute value of the rate of change of morphology – such as the rate of change in circularity – as “morphodynamic speed”.

Protrusions were derived from the binarized image stacks via morphological operations. Each image was first downsized by a factor of 2 in each dimension for the future morphological operations to be computationally feasible. Next, the image was morphologically eroded and dilated with a structuring disk of size 30 pixels in order to remove the protrusions from the cell body. Finally, the binarized protrusion image was found by upsizing the morphologically altered image and subtracting it from the original image.

### E. Optical flow and optical flow alignment

First the actin fluorescence images were jitter corrected using phase cross-correlation [28,29]. Actin optical flow was found by applying iterative Lucas-Kanade [30] successively to each 2-frame combination of images in every fluorescence video. Optical flow confidence is calculated using the spatial Hessian matrix [31] in order to only account for the optical flow vectors that accurately track actin.

The alignment metric was calculated by finding the dot product of each optical flow vector with its surrounding neighbors [32]. To compute this effectively, the optical flow matrix was convolved with a 2D *N* ×*N* gaussian kernel *A* with elements 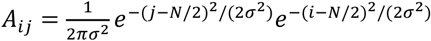with the center (i,j=N/2) set manually to 0. Then, the optical flow alignment was calculated by taking the element-wise inner product of the convolution with the original optical flow matrix. In this study we use *σ* = 1,*N* = 13 in order to capture longer range alignment effects, however the results do not change drastically for values of N between 5 and 31.

The protrusion optical flow alignment (protrusion OF alignment) was found by calculating the average value of the optical flow alignment inside of the protrusion binary image. Previous studies have used different metrics for finding areas of high actin activity that account only for actin fluorescence strength [33], however that does not account for the movement of actin within the cell.

### F. Statistics

Statistical calculations were performed in GraphPad Prism version 10.2 (GraphPad software, CA, US). For comparisons between vehicle control and E2 groups, unpaired Student’s *t*-tests were performed. For all tests, P values < 0.05 were considered statistically significant. Data are represented as mean ± standard error of the mean. Experiments were performed in at least triplicate.

## III. RESULTS

### A. 3D profiling of cell shape dynamics with lattice lightsheet microscopy

We captured cell shape changes, membrane dynamics, and cytoskeleton organization over time using lattice lightsheet microscopy. 12Z cells were cultured, trypsinized, nucleofected with the LifeAct-GFP plasmid for F-actin, and imaged with lattice lightsheet microscopy (Figure 1a). Nucleofection is an electroporation-based method in which an electrical field applied to the cells allows nucleic acids to enter the cytoplasm and nucleus. Cellular dynamics were then visualized with a lattice lightsheet time-lapse microscopy, which illuminates planes of the sample with long thin beams to producie a 3D image with high spatial resolution and minimal photobleaching. To clarify whether the boundary actin dynamics visualized with the LifeAct-GFP probe coincided with the cell membrane, a plasma membrane stain was also added to the cells. Maximum intensity projections of these two channels demonstrate that the boundary of the LifeAct-GFP-expressing cell has a strong overlap with the plasma membrane (Figure 1b), indicating that the actin cortex covers the whole membrane. We caution that the LifeAct-GFP label would not detect actin cortex-free protrusions, such as initial stages of blebs [34]. An example time series of maximum intensity projections from lattice lightsheet imaging of a LifeAct-GFP-expressing 12Z cell with many active protrusions is shown (Figure 1c).

**FIG 1.**
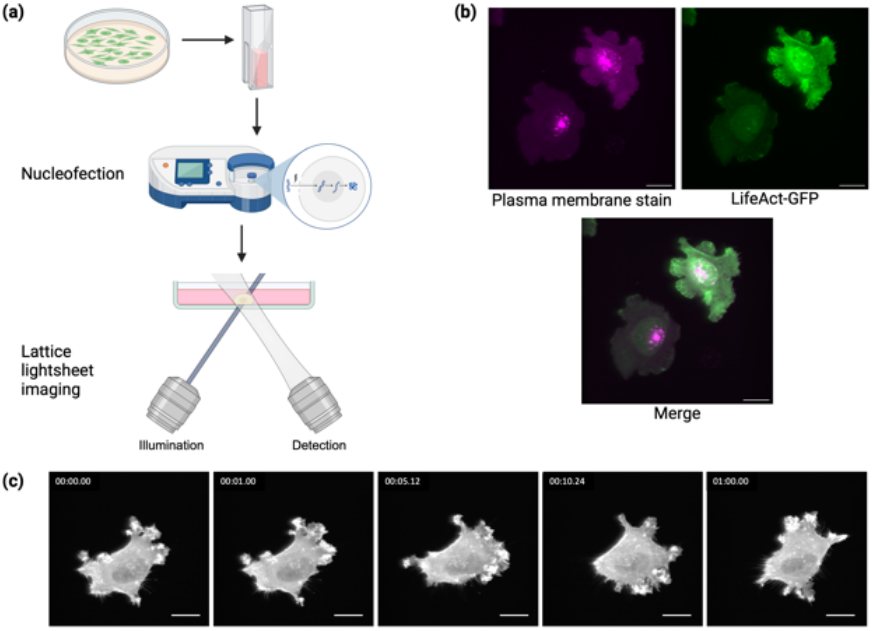
Experimental setup. **(a)** 12Z cells were transfected with a LifeAct-GFP plasmid via nucleofection (an electroporation-based method) and imaged over time using volumetric lattice lightsheet microscopy, which employs long thin beams to illuminate the sample with subcellular resolution. **(b)** A maximum intensity projection showing that the boundary of the actin cytoskeleton (LifeAct-GFP) coincides with the plasma membrane. **(c)** An example output maximum intensity projection from a lattice lightsheet microscopy time-lapse of a cell transfected with LifeAct-GFP with frames from t = 0, 1 min, 5 min, 10 min, and 60 min. Scalebars = 20 µm.

We captured several notable features of the LifeAct-GFP-expressing 12Z cells, indicated with white arrows, at high spatial resolution in 3D from lattice lightsheet microscopy that are visible from isosurfaces (Figure 2). For example, some cells exhibited actin waves pushing against the membrane at the leading edge of the cell, with hair-like protrusions at the trailing edge of the cell (Figure 2a). Several cells exhibited 3D membrane ruffles (Figure 2b and e), filopodia (Figure 2c and d, Figure 2f and g), lamellipodia (Figure 2e and f), and invadopodia-like protrusions (Figure 2d and g). Time series isosurfaces of 12Z cells show membrane ruffle dynamics (Figure 2e), lamellipodia growth and retraction (Figure 2e and f), and protrusion growth and retraction (Figure 2f and g).

**FIG 2.**
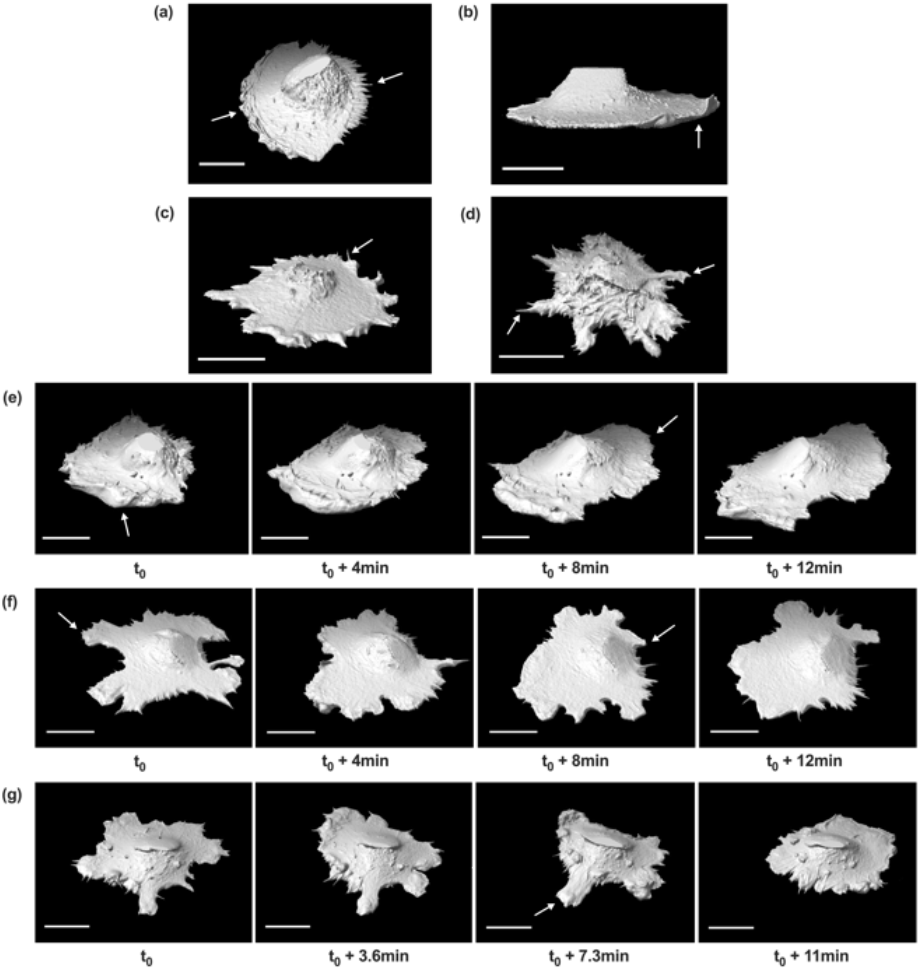
3D isosurfaces of 12Z cells obtained from lattice lightsheet imaging, with some notable cellular features **(a-d)** and representative time series **(e-g)**. (**a)** Some cells exhibited actin waves at the leading edge of the cell and hair-like protrusions at the lagging edge of the cell (indicated by white arrows). Several cells also exhibited membrane ruffling **(b)**, 3D filopodial extensions **(c, d)**, and active protrusions **(d)**. The dynamics of the membrane ruffling and lamellipodia growth were captured **(e)**. Protrusion retraction **(f, g)** and formation **(f)** were also visualized. All scalebars = 20 µm.

### B. E2 treatment alters 12Z cell shape and morphodynamics

To study 12Z morphology and shape dynamics in response to E2 treatment, we analyzed maximum intensity projections of cells transfected with LifeAct-GFP as acquired from lattice lightsheet time-lapse imaging. Cells were treated with vehicle and E2 treatments for 15 minutes or 24 hours and morphological parameters and morphodynamic speeds were averaged over the duration of each experiment (Figure 3). Cells treated for 15 minutes did not exhibit any significant E2-induced changes in circularity or solidity (Figure 3a and 3c, respectively) but did experience more rapid changes in circularity and solidity upon E2 treatment (Figure 3b and 3d, respectively). Of the cells treated for 24 hours, E2 treatment resulted in significantly decreased cell circularity (Figure 3a), solidity (Figure 3c), and rate of change of circularity (Figure 3b) compared to the vehicle control. 24-hr E2 treatment did not significantly alter the rate of change in solidity but a slight downward trend persisted (Figure 3d). In summary, over a short-term period, E2 treatment did not alter cell morphology but did induce more rapid shape fluctuations; over a long-term period, E2 induced clear morphological changes and reduced fluctuations in shape over time (Figure 3).

**FIG 3.**
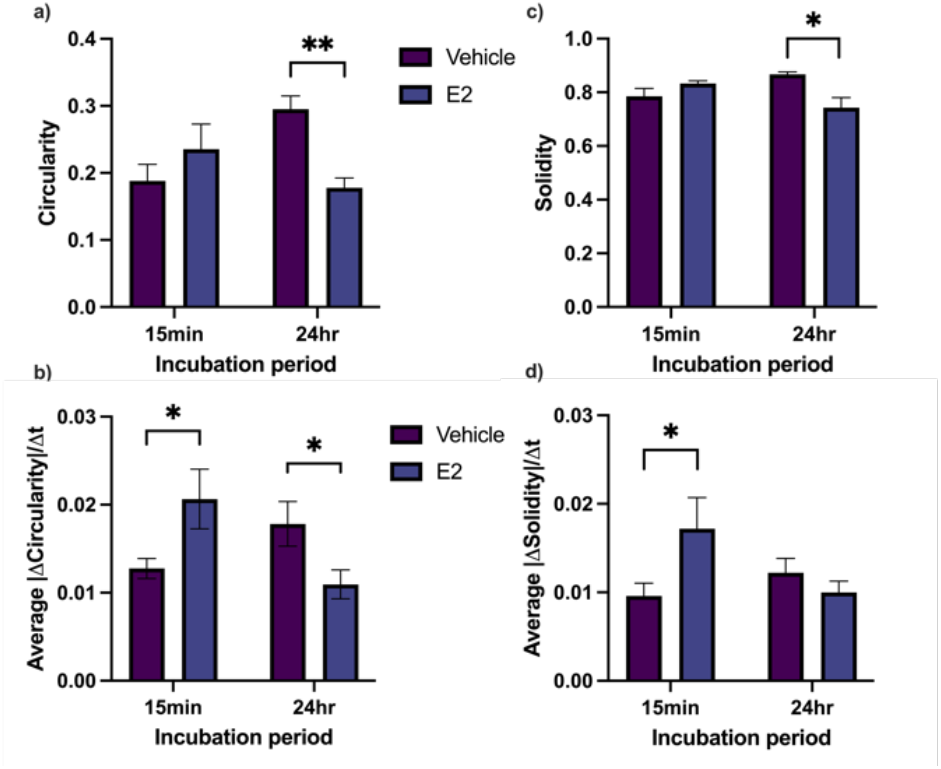
Effect of E2 on 12Z shape and morphodynamics after 15 minutes or 24 hours of treatment incubation. Circularity **(a)**, rate of change in circularity **(b)**, solidity **(c)**, and rate of change in solidity **(d)** over the duration of each time-lapse were extracted from binarized maximum intensity projections, and the means ± SEM are plotted. * P ≤ 0.05, ** P≤ 0.01.

A more in-depth shape-over-time analysis of 12Z cells shows differential behaviors between the vehicle control and E2 groups (Figure 4). Plots of circularity and solidity over time for 15-minute treatment appear relatively disordered (Figure 4a) compared to the noticeable separation between vehicle control and E2 groups for 24 hours of treatment (Figure 4c). Morphological changes normalized to initial values were plotted across different time spans, reported as multiples of 10-second Δt intervals (Figure 4b and 4d). For both 15 minutes (Figure 4b) and 24 hours (Figure 4d) of treatment, short time spans generally resulted in very small morphological changes while longer time spans resulted in larger changes of up to ~15%. Most cells exhibited peak morphological percent changes at 300-fold Δt (equivalent to 50 min), with vehicle-treated cells experiencing larger percent changes than E2-treated cells. Of the cells treated for 15 minutes, vehicle-treated cells tended to increase in circularity and solidity over broad time spans while E2-treated cells tended to decrease in circularity over time (Figure 4b). On the other hand, among cells treated for 24 hours, both the vehicle and E2-treated groups ultimately increased in circularity and solidity over time (Figure 4d).

**FIG 4.**
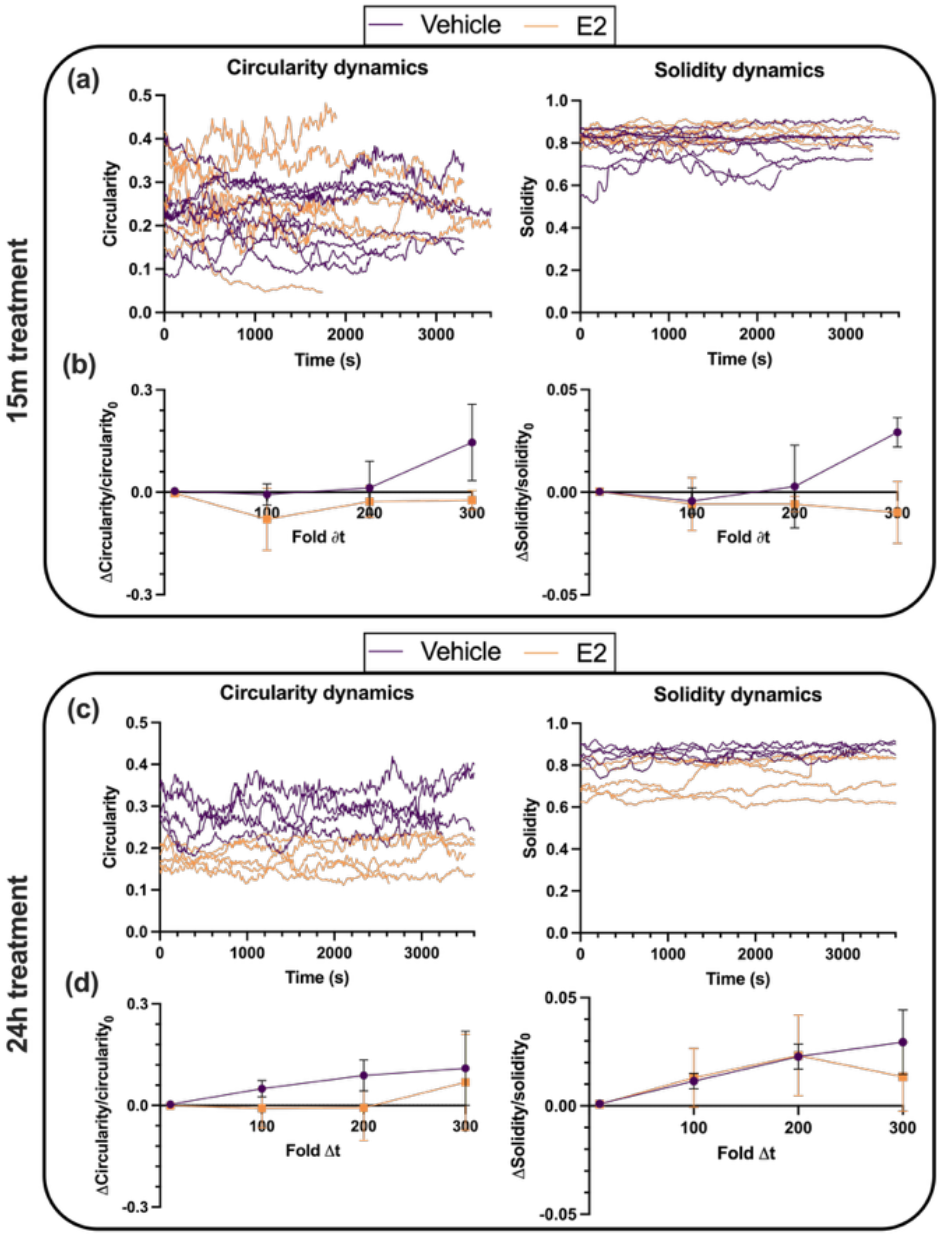
Effect of E2 on 12Z morphodynamics after 15 min (top) or 24 hr (bottom) of treatment incubation. Circularity and solidity were tracked over the duration of each experiment and plotted **(a, c)**, along with the average change in each parameter normalized to initial value over time intervals of varying lengths **(b, d)**. The base time step Δt = 10 seconds.

### C. E2 treatment increases membrane protrusions in 12Z cells after 24 hours

To study the effect of E2 on 12Z membrane protrusions, protrusion area and cell area were quantified from fluorescence maximum intensity projections. We automatically segmented the protrusions using morphological operations (see Materials and methods) and two broad categories of cell protrusion dynamics were seen. The first category observed was invadopodia-like dynamics (Figure 5a) which are characterized by large, non-uniform protrusions with no front of the cell clearly demarcated. Invadopodia are commonly observed in cancer cells as a means for metastasis and migration [22], however here we use the term due to visual similarities between the protrusions in our cells and true invadopodia. The other category observed was lamellipodia-like protrusions (Figure 5b), which we classify as cells having broad, flat protrusions focused mainly at the front of the cell. These protrusions tend to be much smaller than the size of the cell and are largely present in unidirectional guidance [35].

**FIG 5.**
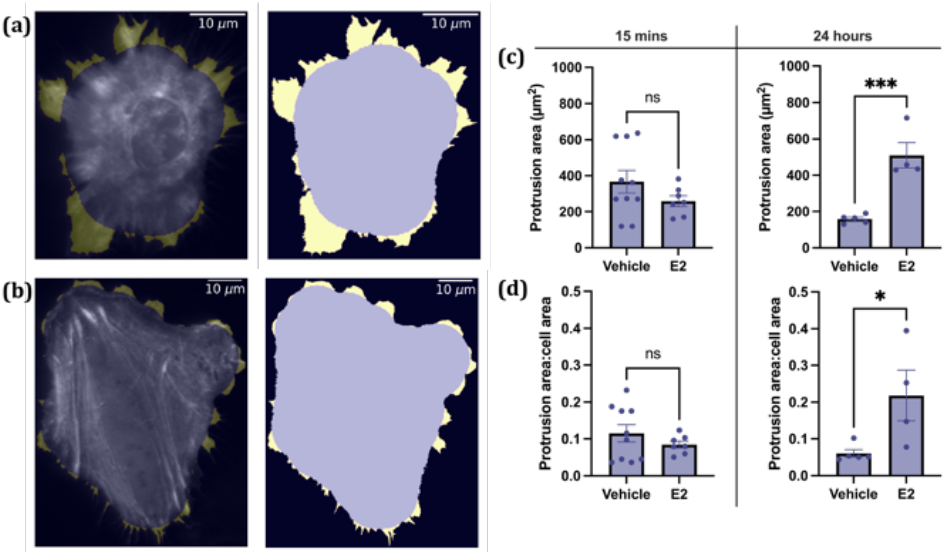
Effect of E2 on cellular protrusions. **(a)** A cell with invadopodia-like dynamics, which are characterized by large protrusions on all sides of the cell. The first image of the series shows cell protrusions in yellow overlaid onto the actin fluorescence image. The second image shows the binarized mask of the cell body (purple) and protrusions (yellow). **(b)** A cell with lamellipodia-like dynamics, which are characterized by broad short protrusions. **(c)** Protrusion area in µm^2^ and **(d)** protrusion area:cell area ratio were extracted from binarized images and the means ± SEM are plotted. * P ≤ 0.05, *** P≤ 0.001.

To determine how E2 treatment affected protrusion dynamics we found the protrusion area for all cells. E2 treated cells have no change in protrusion area compared to vehicle control cells after 15 minutes (Fig 5c, left panel), but significantly larger protrusions than vehicle control cells after 24 hours of treatment (Figure 5c, right panel). We accounted for variability in cell size by looking at the ratio of protrusion area to cell body area. Figure 5d shows that after 15 minutes of E2 treatment (left panel) the ratio of protrusion area:cell area does not change compared to the vehicle control cells. Again, this difference is significant for the cells treated for 24 hours (Figure 5d, left panel), indicating that result holds across cell sizes.

### D. E2 treatment decreases actin optical flow alignment in 12Z cells after 24 hours

To this point we have focused mainly on cell morphodynamics; however, we have fluorescently tagged actin in the cell to investigate the impact of E2 treatment on the dynamic shift in the location of the F-actin scaffolding. Dynamics may be driven by a combination of polymerization, depolymerization, myosin contraction, or cytosolic flow. To quantify actin dynamics we use optical flow [32] to find the speed that the actin is moving within the cells. The optical flow is a 2D vector field (Figure 6a) that gives the shift in location of the actin scaffolding as well as an approximation of the speed with which it moves. Areas of the cell with higher actin optical flow have faster change in their scaffolding at that point. However, there was no significant difference in the actin optical flow distributions between the E2-treated cells and the vehicle control cells.

**FIG 6.**
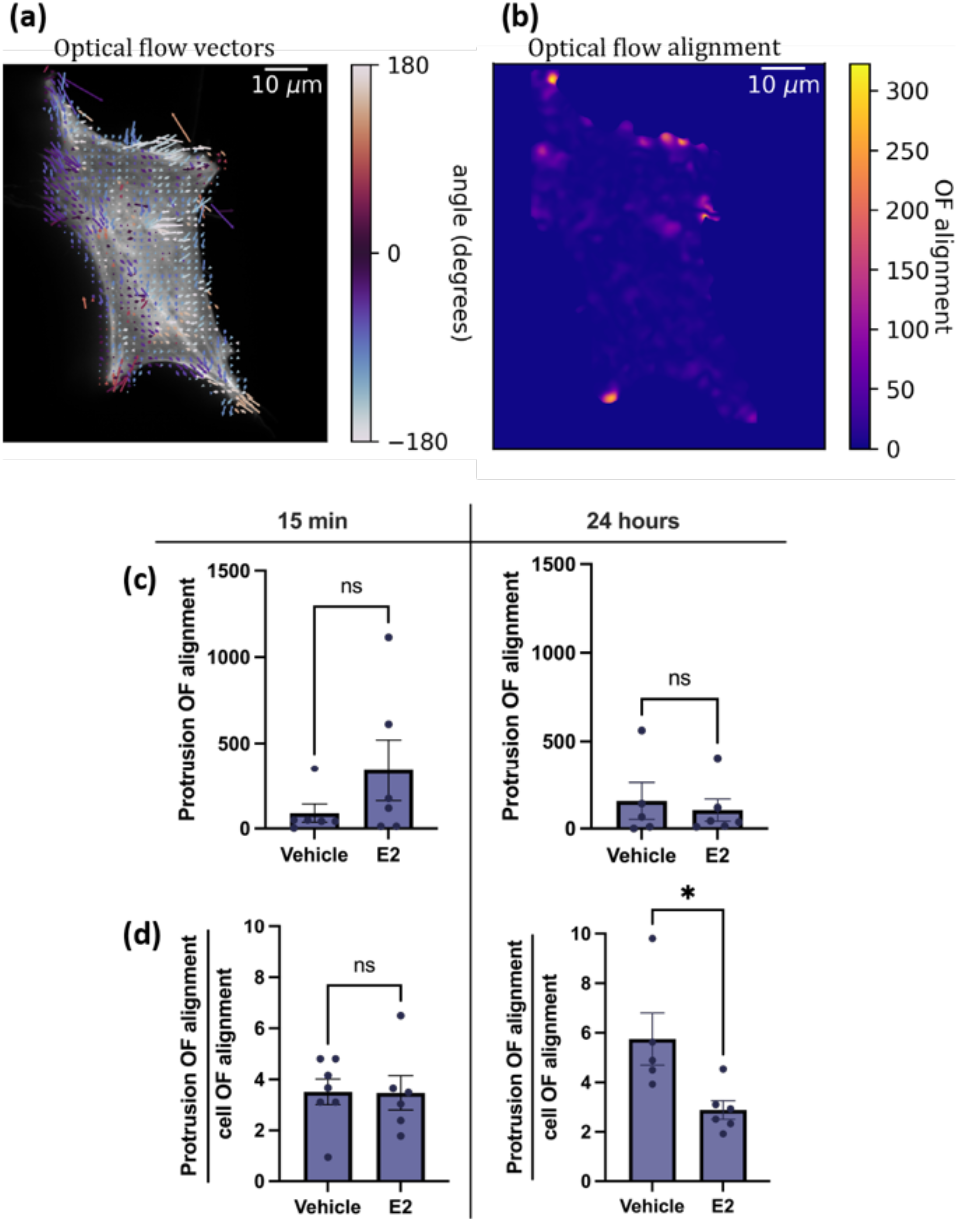
Impact of E2 on actin optical flow alignment in 12Z cells. **(a)** Optical flow (OF) was calculated from actin fluorescence for all timepoints to determine how the actin is moving within the video. The optical flow vectors are displayed on top of one of the corresponding actin fluorescence images. **(b)** The actin optical flow alignment (OF alignment) is displayed for the cell in **(a)**. Higher values of optical flow alignment mean that the optical flow in that region is pointing in the same direction (see Materials and methods). Using the binarized masks from Fig. 5 the average optical flow alignment was found inside the protrusions and the cell body. The mean protrusion optical flow alignment **(c)** and mean ratio of protrusion optical flow alignment to cell body optical flow alignment **(d)** are plotted ± SEM. * P ≤ 0.05, ** P≤ 0.01.

In order to quantify how the actin was moving collectively within the cell, we calculated the alignment of the optical flow vectors (Figure 6b). Regions of high alignment indicate regions where coordinated shifts in the cellular scaffolding occur, for instance an actin polymerization wave that travels in one direction. The optical flow alignment was calculated by taking the spatial average of the dot product of each optical flow vector with its neighbors (see Material and methods). At any given point, higher alignment means the optical flow vectors around that point have similar directions. Lower values of alignment either mean that the optical flow was very small in that region or that the vectors are unaligned. Since coordinated shifts in the scaffolding are needed to generate protrusions, we expected the alignment to be higher in protrusions than in the cell body. Therefore, we calculated the average alignment in both the cell body and the protrusions separately.

Figure 6c shows no significant difference in the average protrusion optical flow alignment; the protrusions have similar levels of actin alignment in both treatments and incubation times. However, if we consider the ratio of protrusion optical flow alignment:cell body optical flow alignment we see that the ratio is significantly lower for the E2-treated cells compared to the vehicle control in the 24-hour group, but not the 15-minute group (Figure 6d). This means that the E2 treatment, after 24 hours, makes the cell’s actin significantly less aligned in the protrusions compared to the bulk of the cell. With the E2 treatment, the actin in the protrusions becomes more disordered compared to the actin in the bulk.

## IV. DISCUSSION

We used lattice lightsheet microscopy to capture high-resolution cell dynamics, enabling an unprecedented visualization of protrusive activity and actin polymerization dynamics in live endometriotic cells. We showed that this technique can be used to visualize a variety of 3D cytoskeletal structures, including those associated with ruffles, filopodia, lamellipodia, and invadopodia-like protrusions over time.

This study aimed to elucidate the effect of E2 on actin and shape dynamics using the 12Z human endometriotic epithelial cell line. We demonstrated that 24-hour E2 treatment significantly reduced average cell circularity and solidity compared to the vehicle control –– changes indicative of a transition to a more invasive phenotype as cells adopt a more elongated, protrusive shape. In the short term (15 minutes), E2 did not alter cell morphology but did induce more rapid morphodynamic changes. This contrasts with 24-hour E2 treatment, which resulted in slower morphodynamic changes. These results suggest that E2 may stimulate rapid, transient fluctuations in the short-term –– potentially due to rapid, non-classical estrogen signaling pathways –– and cause a more stable but invasive morphological state in the long-term –– a time scale more on par with classical estrogen signaling that directly regulates transcription [36].

In line with the results in Figure 3, detailed shape analysis over time revealed that although E2-treated cells are highly morphodynamic after 15 minutes, clear, stable morphological shifts are only seen after 24 hours. Both vehicle and E2 groups experienced shape changes that accumulated over time. Peak morphological changes (up to ~15%) occurred over a window of 15-50 minutes, which may reflect the time scale of 12Z membrane protrusion turnover. After 24 hours, vehicle-treated cells generally exhibited larger fluctuations in circularity compared to E2-treated cells, which settled into a more stable but invasive morphology. Given that E2 is generally known to promote cell invasiveness in several diseases [1,7,11,12], this result matches our expectation. Future studies could clarify the mechanisms underlying E2-mediated morphodynamic states in the 12Zs; for example, investigating the distinct, time-dependent contributions of classical and non-classical estrogen signaling pathways to cytoskeletal remodeling would elucidate the impact of E2 on cell shape over time.

We discovered that 24-hour E2 treatment leads to different protrusion dynamics as well as modified actin dynamics within those protrusions compared to vehicle control cells. E2-treated cells generally displayed invadopodia-like dynamics, with high protrusion areas and high ratio of protrusion area:cell body area. These higher area protrusions tended to jut out of the cell in every direction. Large protrusions in non-polarized cells tend to be a marker of invasive cells [37,38]. This suggests that the E2-treated cells are primed to be more invasive than the vehicle control cells, because they are forming larger, non-uniform protrusions that are reminiscent of invadopodia. This effect appears only after 24 hours of treatment, not after 15 minutes, suggesting that E2-induced invadopodia-like protrusions are not formed as a result of rapid signaling. Future work could investigate the expression of invadopodial proteins, such as cortactin or MT1-MMP, in 12Z protrusions and determine whether E2 treatment enhances their proteolytic activity. This would clarify mechanisms driving E2-mediated invasion in benign and malignant estrogen-dependent conditions.

Within the membrane protrusions, the actin behaved differently in E2-treated cells. After 24 hours, the ratio of protrusion actin alignment:cell body actin alignment was significantly higher in vehicle-treated cells than E2-treated cells, indicating that the actin was more well-aligned in the vehicle control. When E2 was added to the cells, it appeared to make the actin in the protrusions more disordered. This suggests that actin polymerization is less coordinated in the E2-treated cells, which would be beneficial in sensing non-uniform local topography. This exploratory, protrusive state could support cell invasion in complex 3D environments where an adaptable cell shape is advantageous. Prior studies show that estradiol activates both RhoA and p21-activated kinase (PAK) [15], which are essential regulators for the formation of lamellipodia and pseudopodia [39,40]. Our results suggest that this change in RhoA and PAK may result in more disordered actin polymerization over a 24-hour time scale.

Some limitations of this study should be noted. LifeAct-GFP, a widely used F-actin probe, has been previously shown to influence actin dynamics in other cell types [41–43] and has been unable to stain actin-based filopodia within chick embryo mesenchymal cells [44]. LifeAct-GFP also has a relatively high affinity to G-actin [45] which generates background fluorescence. Although our imaging validates LifeAct-GFP localization with plasma membrane staining, other methods of fluorescently labeling F-actin should be tested and compared with our results. Additionally, many cells that were imaged experienced a reduction in cell area over time, especially after about one hour of imaging, which could be due to some degree of photosensitivity. Future studies that image other cell types using lattice lightsheet microscopy will provide more context on the relative photosensitivity of the 12Z cells. Finally, we were limited in our ability to quantify the 3D dynamics of actin due to computational constraints. However, we demonstrated that the maximum intensity projection of the 3D data can be effectively analyzed for morphodynamics and actin dynamics. Future advances in computational analysis pipelines and hardware will allow us to gain further insight on cytoskeletal remodeling in 3D.

Together, our findings suggest that estradiol alters cytoskeletal plasticity and shape dynamics in endometriotic epithelial cells, potentially priming them for increased invasiveness. These insights lay the groundwork for future studies exploring the molecular signaling pathways and mechanical feedback loops that underlie hormone-driven motility in estrogen-responsive processes.

## ACKNOWLEDGMENTS

The authors acknowledge funding from an NIGMS MIRA #R35GM142838 (to KMS), from an NSF grant PHY-2014151 (to WL and CH), and from the Clark Doctoral Fellowship (to SB). We thank the Biosciences Core Imaging Incubator at the University of Maryland for providing access to the Zeiss LLS7 microscope. Schematics in Figure 1 were created with Biorender.com.

## AUTHOR CONTRIBUTIONS

SB performed sample preparation, data collection, and data analysis. CH performed data collection, code development, and data analysis. SB and CH drafted the manuscript together. KMS and WL provided guidance on the experiments and analysis, manuscript structure, and figures. All authors reviewed the final manuscript.

